# Factors influencing prevention of mother to child HIV transmission service utilization among HIV positive women in Amhara National Regional State, Ethiopia: A thematic content analysis

**DOI:** 10.1101/613752

**Authors:** Zebideru Zewdie Abebe, Mezgebu Yitayal Mengistu, Yigzaw Kebede Gete, Abebaw Gebeyehu Worku

**Author notes:** Corresponding author: Zebideru Zewdie, Ethiopian Federal Ministry of Health, Addis Ababa, Ethiopia, P.O. Box: 1234.

## Abstract

**Introduction:** Mother to child transmission (MTCT) of HIV is the major source of HIV infection among children under the age of 15 years. Prevention of mother to child transmission (PMTCT) service has been an important strategy in preventing HIV infections in infants. However, improving PMTCT service uptake and continuum of care still remains a significant impediment in the Amhara Region of Ethiopia. The aim of this study was to explore factors that may hinder and promote PMTCT service utilization among HIV positive women.

**Methods:** Phenomenological study design was used. Three focus group discussions (FGDs) with HIV positive women and five in-depth interviews with health care workers were conducted from the selected health institutions. Data analysis was conducted using thematic content analysis. ATLAS/ti version 7.5.16 software was used to assist in coding and analysis of the qualitative data.

**Results:** The findings of the study revealed that there are a range of factors that influence HIV positive women on PMTCT service utilization. Stigma, discrimination, limited knowledge on HIV, lack of partner and family support, and not getting friendly service from health facility were factors that might hinder PMTCT service utilization. The experiences of HIV positive mothers revealed that continuous partner support, previous history of getting HIV free child, good health worker support, and advice by mother support group were factors that promote utilization of PMTCT service.

**Conclusions:** Strengthening community awareness on HIV, engaging male involvement in PMTCT care and getting friendly service were the key determinants for the better PMTCT service utilization.

## Introduction

Mother to child transmission (MTCT) of HIV is the major source of HIV infection among children under the age of 15 years. Without intervention, the risk of MTCT ranges from 20% to 45%. With specific interventions the risk of MTCT in breastfeeding and non-breastfeeding populations, can be reduced to less than 2% and 5% respectively [1]. Ethiopia is among the 21 prioritized countries with the highest number of pregnant women living with HIV globally in 2014 [2].

Women of childbearing age account for more than half of the world’s HIV cases. Unintended pregnancies are high among HIV-positive women [3]. Added to this, over 90% of HIV infections in children are acquired through MTCT. Studies indicated that without appropriate PMTCT interventions about 50% of those infants born with HIV will die before their second birthdays [4]. PMTCT is a highly effective intervention and has huge potential to improve both maternal and child health [5]. According to the WHO guideline there are four comprehensive approaches or prongs to PMTCT programs: preventing new HIV infections among women of childbearing age; preventing unintended pregnancies among women living with HIV; preventing HIV transmission from a woman living with HIV to her baby; and providing appropriate treatment, care and support to mothers living with HIV and their children and families [6, 7]. Moreover, the Sustainable Development Goals (SDGs) place heightened emphasis on PMTCT in the context of better health for mothers and their children [1].

According to 2015 UNAIDS progress report, the percentage of pregnant women living with HIV who were receiving antiretroviral therapy increased from 64% in 2013 to 73% in 2014 in Ethiopia [8]. The report also revealed that the incidence of new HIV infections in children reduced by 65% since 2009. However, the challenges of maintaining women on antiretroviral medicines throughout the breastfeeding period still persist. In addition, only 25% of HIV-exposed infants received early infant diagnosis, and 22% of children (aged 0–14 years) living with HIV received antiretroviral therapy in 2014 [8]. The involvement of male in PMTCT program is also low. In addition the time for mothers to bring infant to conduct infant diagnosis were delayed. In 2014, the percentage of mother to child HIV transmission (MTCT) is high (18%) in Ethiopia [8]. Even though implementation of option B+ strategy (test and treat) started in 2013, the status PMTCT service utilization in the PMTCT care cascade are still low in Amhara National Regional State, Ethiopia.

PMTCT service utilization was influenced by different factors. Studies revealed that factors associated with the uptake of PMTCT services are knowledge and attitudes of mothers on HIV, stigma and discrimination, male partner involvement, disclosure of HIV status, quality of the service, health workers approach, access to and availability of PMTCT services, and educational status of the women [9-12].

Prevention of communicable disease including HIV/AIDS is the Ethiopian government top priority. Availability of PMTCT and ART service will lead to reduction of HIV infection to reproductive health women and prevent pediatric HIV and improve child survival. However, improving PMTCT service uptake and continuum to care still remains a significant challenge in Amhara Region, Ethiopia. Understanding barriers and facilitators of PMTCT service utilization is used to improve service uptake and ensure optimal outcomes for women and their infants. Therefore this study was conducted with the aim to explore factors that may hinder and promote PMTCT service utilization among HIV positive women in Amhara Regional State, Ethiopia.

## Methods

### Study design and setting

Phenomenological study design was employed to explore factors that may hinder and promote PMTCT service utilization among HIV positive women in Amhara Regional State, Ethiopia. The study was conducted in three health centers of Amhara region which provide PMTCT service. Amhara region is located in the northwestern part of Ethiopia. The total population of Amhara region is estimated to be about 20 million with mean annual growth rate of 1.7% [13] in which female population accounts for 49.6% of the population and there is an estimated 4,345,893 women of reproductive age (15-49 years). The estimated HIV positive women in need of PMTCT services in the region in 2017 were 8,720 [14]. The region has 5 referral hospitals, 3 zonal hospitals and 32 district hospitals. The region also has 832 health centers and 3336 health posts [15].

### Study population and sampling

The study population of this study was all HIV-positive women who were on follow up visit at PMTCT in selected health facilities of Amhara region. In addition, health workers who were providing the PMTCT service at selected hospitals and health centers were included in the study. However, all HIV-positive women who were seriously sick were excluded from the study.

Three focus group discussions (FGDs) with HIV positive women and five in-depth interviews with health workers providing PMTCT service were conducted at health centers of Amhara Region, Ethiopia, in 2018. HIV positive women who came for PMTCT service in the selected health facility were selected for FGD purposively. In addition to HIV positive women, health workers rendering PMTCT services at selected health facilities were included in the in-depth interview.

### Data collection procedures

Focus group discussion with HIV positive women attending PMTCT unit was conducted to explore factors that may hinder and promote PMTCT service utilization. The data collection instruments were focus group discussion and in-depth interview. The focus group discussion was conducted after taking informed consent from the participants. The number of women per each focus group varied, the minimum being seven and the maximum 10. The FGD were carried out in quiet place in the health facility and the in-depth interview was also carried out privately and without interruption. The discussions were in local language which is Amharic and tape recorded. The duration of the discussions ranges from 45 min to one hour. Two facilitators (one moderator and one note taker) were guiding the focus group discussions.

The focus group discussion was began with questions regarding general information about HIV/AIDS and PMTCT, and then the focus was directed toward an in-depth discussion of factors associated with the utilization of PMTCT services. The FGD guide includes information on PMTCT service intervention like ARV uptake, utilization of dual contraceptive, early infant diagnosis, Neverapin prophylaxis and exclusive breast feeding for their HIV exposed infants for PMTCT.

### Data management and analysis

Data analysis for both the in-depth interviews and the FGD was conducted using thematic content analysis. Qualitative data were transcribed and translated into English, and imported into Atlas-ti version 7.5.16 software for coding and analysis. Data was organized along the main themes of the study using existing themes in the interview guide and emerging themes, after reading through the transcripts several times. Related quotations were merged into categories/subtheme, which later formed the themes. Focus group discussions had facilitated an in-depth exploration of factors that may hinder and promote PMTCT service utilization.

### Ethical Considerations

The study was conducted after obtaining ethical clearance from Ethical Review Board of the University of Gondar. Support letters from the Regional Health Bureau and Zonal Health Department were secured. Written informed consent was obtained from all research participants after explaining the purpose of the study. The aim, benefits and disadvantages of participation in the study, and the time taken for FGD and In-depth interview were explained to study participants. Confidentiality, anonymity and privacy were maintained by assigning codes to each participant. Finally the participants were approached to take part in the study and consent was signed if they decided to participate.

## Results

### Socio-demographic description of participants

A total of 26 HIV-positive women who had follow-up in PMTCT clinic participated in the FGDs. Of the participants, 25 (96.2%) were married and 22 (84.6%) of participants were on ART for more than one year and all participants were currently on ART. The mean age of study participants who participated in the FGDs was 30.9 years, ranging from 20 to 40 years (Table1). In addition five health care workers working in PMTCT unit participated in the in-depth interview.

**Table 1:** Demographic and PMTCT service utilization characteristics of study participants in selected health facilities, Amhara Regional State, Ethiopia, 2018

| Demographic characteristics | Number (n= 26 ) | Percentage (%) |
| --- | --- | --- |
| **Age range (years)** |  |  |
| 20 - 24 | 2 | 7.7 |
| 25_29 | 7 | 26.9 |
| 30_34 | 9 | 34.6 |
| 35_39 | 7 | 26.9 |
| 40+ | 1 | 3.9 |
| **Time since diagnosis** |  |  |
| < 1 year | 3 | 11.5 |
| 1 year | 1 | 3.8 |
| 2- 5 years | 6 | 23.1 |
| 6- 9 years | 10 | 38.5 |
| 10 and above years | 6 | 23.1 |
| **Using ARV at time of interview** |  |  |
| Yes | 26 | 100.0 |
| No | 0 | 0.0 |
| **Marital status** |  |  |
| Married | 25 | 96.2 |
| Single | 1 | 3.8 |

Similar quotations were categorized under sub-themes/categories and these categories further merged to themes. A total of 34 sub-categories, eight categories/sub-themes and three themes/families emerged from qualitative data analysis. The sub-categories were ranked here in order of the frequency with which they were provided by respondents. Therefore, the findings of this study were presented with three themes namely awareness about HIV risk, facilitators, and barriers to the PMTCT service utilization. Verbatim quotations from the study participants were mentioned without any attempts to correct grammatical errors. Detail discussions were made on the three themes based on the objective of the study (Table 2).

**Table 2:** Factors that may hinder and promote PMTCT service utilization among HIV positive women in Amhara Regional State, Ethiopia, 2018.

| Theme | Category | Sub-category |
| --- | --- | --- |
| Awareness about HIV risk | Awareness on HIV transmission and PMTCT | Awareness about HIV transmission |
|  |  | Awareness about PMTCT of HIV |
|  |  | Awareness about MTCT of HIV |
|  | Sources of information for PMTCT | Media (Radio, Television) |
|  |  | Health workers |
|  |  | During ANC follow up |
|  |  | Health education |
| Barriers to PMTCT service utilization | Community related factors | Stigma |
|  |  | Discrimination |
|  |  | Lack of awareness |
|  | Individual and family related factors | Fear of divorce |
|  |  | Lack of partner support |
|  |  | Lack of family support |
|  |  | Lack of time |
|  |  | Nutritional problem |
|  |  | Religious beliefs |
|  |  | Stigma |
|  |  | Discrimination |
|  |  | Lack of awareness |
|  | Health institution related factors | Cost of the drugs like cotrimoxazole |
|  |  | Not getting friendly service from health facility |
|  |  | Problem on giving the service from the health care providers |
|  |  | Long waiting time to get service |
| Facilitators to PMTCT service utilization | Community related factor | Good willingness for disclosure of HIV status |
|  | Individual and family related factors | Continuous partner support |
|  |  | Willingness to disclosure of HIV status |
|  |  | Good support from family |
|  |  | Previous history of getting HIV free child |
|  |  | Feeling Healthy |
|  |  | Good awareness on HIV transmission |
|  | Health institution related factors | Good Health worker support |
|  |  | Awareness creation |
|  |  | Availability of drugs |
|  |  | Discussion during coffee ceremony with mother support group (MSG) |
|  |  | Availing other supplementary support |

### Awareness on HIV risk

This theme merged from sub-theme related to the awareness about HIV transmission and PMTCT, and sources of information about PMTCT. Most of the study participants knew about the HIV transmission methods. Most of the participants mentioned three to four transmission methods like unsafe sex, during blood transfusion, mother to child transmission and using sharp objects.

A 27 years old HIV positive woman explained that the main HIV transmission methods as “the main are three, blood contact, mother to child transmission in three ways during pregnancy, labor and breast feeding time, unsafe sex. In addition HIV can be transmitted by contact and when giving care to AIDS patients taking prophylaxis is important”. Another 30 years HIV positive woman participant added on the methods of HIV transmission; “unsafe sexual relation, using sharp objects together and mother to child transmission”. But still there is a knowledge gap on HIV transmission method. For example, 24 years old married HIV positive women explained that “I have infected by knife during meat cutting in my friend’s ceremony (Digis)”.

Most of the study participants had awareness about mother to child transmission of HIV. Those who mentioned MTCT indicated that it could occur during pregnancy, during delivery and through breastfeeding. Most women had a good understanding on PMTCT and awareness that interventions to prevent such transmission were available. But few participants mentioned only ART to mother and NVP prophylaxis to infant, no one mention other interventions.

The main sources of information about PMTCT mentioned by the participants were health workers, radio/television and health education during antenatal visit. Health workers and radio/television were the frequently mentioned sources of information.

### Barriers to PMTCT service utilization

Participants mentioned different hindering factors to PMTCT service utilization during FGDs. The perceived barriers described by participants categorized in to sub themes like community factors (stigma, discrimination, limited knowledge on HIV); individual and family related factors (stigma, discrimination, no partner support, fear of divorce, religious beliefs, lack of family support, lack of time, nutritional problem, lack of awareness); and health institution factors (cost of the drugs, problems on provider side, not getting friendly service from health facilities, taking time to get service) (Table 2). The detail in each subtheme/category presented below.

According to the health workers responses, the main reasons for women not taking PMTCT service were disclosure of the result, fear of partner, drug side effect, drug supply interruption for example Cotrimoxazole, lack of attention by the health care workers and work load.

### Community Factors

This category refers to community factors for PMTCT service utilization. The most frequently mentioned factors related to the community were stigma, discrimination and lack of awareness about PMTCT by the community.

#### Stigma

The influence of their peers, family and community on women can be tough if her HIV status is known. This was expressed by a 33 years old HIV positive FGD participant as **“**No one sit on a place where an HIV person sits. That is why I hide my status from my family and now I took the drugs for 8 years”.

#### Discrimination

Most of the participants recognized that HIV/AIDS stigma was still prevalent within the community and women living with HIV/AIDS were discriminated due to their HIV status. For example HIV positive mothers afraid their house renter and not taking drug regularly. This was expressed by a 33 years old HIV positive FGD participant as **“**I was fired from rented house in the second day of my delivery. The house renter asks me why you give syrup for newborn and I told her that it is HIV prophylaxis then she told me to leave the house immediately”. Another 35 years HIV positive woman participant added on the barriers for not taking the drugs continuously; “When a guest comes we may discontinue taking the drug because we afraid of them”

### Individual and family related factors

This category refers to individual and family related factors for PMTCT service utilization. Frequently mentioned individual and family related factors identified during FGD were fear of divorce, lack of family support, lack of partner support, and lack of time. Nutritional problem, religious beliefs, stigma, discrimination, and lack of awareness on PMTCT were also noted, but less frequently.

#### Fear of divorce

Fear of divorce was the most frequently cited barrier for not taking the PMTCT service and continuing treatment. A 27 years HIV positive participant said that “most of HIV positive men are not transparent to their partner; the husband takes drugs without the knowledge of his partner but mothers afraid not to disclose their status due to afraid of divorce”. Similarly a 27 years old HIV positive participant further explained this issue as; “I know one mother her husband fired out from her house and get HIV positive child due to disagreement and not taking proper care”. A 24 years FGD participant also echoed “She discontinued the drug due to afraid of her husband and child”. Another 30 years old participant also stressed on the importance of HIV status disclosure to partner. She said that “disclosure of the HIV status is necessary for all people, I have divorced from my husband due to lack of transparency”. Health workers rendering the PMTCT services also explained that “fear of partner is the main reason for HIV positive women for not disclosing their HIV status and not taking PMTCT services”.

#### Lack of family and partner support

Lack of family support was the most frequently cited barrier for not taking the PMTCT service. This is expressed by a 22 years old participant as “afraid of other people is the main reason for not taking the drug properly, for example my family may stigmatize me like if I wear their cloth they perceive that HIV can be transmitted to them and also they did not drink with the same glass with me. If I was sick I get counseling from health workers not from my family”. We found evidence of other barriers like limited knowledge about PMTCT by the community, lack of time, nutritional problem and limited knowledge of HIV but less commonly mentioned.

### Health care service factors

This category refers to health care service factors for PMTCT service utilization. Health care related factors such as high cost of the drug, not getting friendly service from health facility and long waiting time were frequently mentioned factors for not utilizing PMTCT service. This was expressed by a 24 years HIV positive woman as “it is costly and difficult to buy infant syrup by the poor”. Another participant, a 39 years old HIV positive women also explained that “The service provided to us is good but they order syrup to child to buy from outside, it is costly and difficult to buy for the poor”. A 33 years old participant also reiterated that “Giving child syrup free of payment because most of the mothers have no enough income”. Health workers rendering the PMTCT service also explained that “drug supply interruption for example Cotrimoxazole shortage was a problem for PMTCT service utilization”.

Getting friendly services from health facility is important to improve service uptake. This is expressed by a 30 years HIV positive participant explained her concern about not getting good service from health facility as “in my experience getting proper services is difficult, I stayed for 3 days at hospital before delivery as rapture of membrane and leaking of amniotic fluid without any support. I born my child at hospital, but the fetus was dead”. A 35 years old HIV positive participant echoed that “if you told to health workers at hospital that you are HIV positive, you cannot get proper services”. Similarly health workers providing PMTCT service also explained that “trained health care professionals have not given attention to the service due to work load”.

### Facilitators for PMTCT service utilization

Participants mentioned different promoting factors to PMTCT service utilization during FGDs. These promoting factors categorized in to sub themes like community factors (good willingness to disclose HIV status); individual and family related factors (continuous partner support, willingness to disclose HIV status, good support from family, previous history of getting HIV free child, feeling healthy, good awareness about HIV transmission); and health institution factors (good health worker support, awareness creation, availability of drugs, discussion during coffee ceremony with mother support group, and availing other supplementary support) (Table2). The detail in each subtheme/category presented below. Health workers also mentioned different factors that promote PMTCT service utilization during in-depth interview. The most frequently mentioned promoting factors mentioned by health workers were orientation to all clinical staffs, uninterrupted drug supply, strengthen mother support group counseling, availing coffee ceremony budget, and arranging separate room for counseling.

### Community Factors

The most frequently mentioned promoting factors related to the community was good willingness to disclosure of HIV status. This was expressed by a 40 years HIV positive woman as “giving health education about healthy living, health care and PMTCT was important”.

### Individual and family related factors

The frequently mentioned promoting factors related to individual and family were partner support, disclosure of HIV status, good support from family, previous history of getting HIV free child, healthy feeling, and good awareness on HIV transmission. Continuous partner support to the women for those who had disclosed their HIV status was reported to the most frequently mentioned promoting factors for PMTCT service utilization. This support can be like remind to take their drugs and counseling. This was expressed by a 33 years participant as “my partner leaves away from me but he remembers me to take the drug by telephone”. A 36 years old HIV positive participant reiterated that “I am a daily laborer and have two children; my child supports me to take my drug properly”. Another participant 27 years old HIV positive women also explained that “Having HIV free child gives happiness to the mother and this promote for regularly follow of service to have HIV free child; When I know my child is free I was the happiest person”.

### Health care service factors

One of the major facilitators for PMTCT service utilization and retention in care was good support services provided by health workers. Awareness creation on the service provided, discussion on coffee ceremony by mother support group and availability of drugs were also frequently mentioned promoters for PMTCT service utilization. Availing other supplementary support like nutritional support was also promoting PMTCT service utilization. This was expressed by a 28 years HIV positive woman as “All HIV positive people feel healthy and no one sick at this time due to this they are motivated to take drugs properly”. Another participant, 30 years old HIV positive women also explained that “discussion during coffee ceremony by mother support group is important to share experiences”. A 31 years old HIV positive participant reiterated that “Health workers counseling and giving service with respect and patience”. Another participant (Health workers working at PMTCT clinic) also explained that “strengthening mother support group counseling and availing coffee ceremony budget are important to improve PMTCT service utilization”. A 39 years old HIV positive woman said that “Previously our friends were given different support like blanket and oil”.

## Discussion

This study identified a range of factors that restrict HIV positive women from participating in PMTCT follow-up across the PMTCT care cascade. The most frequently mentioned factors related to the community were stigma, discrimination and lack of awareness about PMTCT by the community. Most of the participants recognized that HIV/AIDS stigma was still prevalent within the community and women living with HIV/AIDS were discriminated due to their HIV status. The findings described in this study were similar to study conducted in Kenya that showed stigma and discrimination are reported as the main hindrances to PMTCT service uptake [16]. Other studies have also reported similar findings [17, 18]. Being known by the community as to be HIV infected, and fear of stigma can impact on a woman’s decision on whether or not to get care for herself and her infant.

Frequently mentioned individual and family related factors identified in this study were fear of divorce, lack of family support, lack of partner support and lack of time. Like our research, other studies found that non-disclosure of HIV status, fear of divorce, fear of HIV stigma, and lack of partners support were some of the most common barriers to PMTCT service utilization [16, 19-26]. Another study in Sudan [27], and in Malawi [28] also revealed that women are not likely to disclose their HIV status due to stigma. This indicated that family support is important to get health service but stigma and discrimination affect family support.

This study also revealed that health care related factors such as cost of the drug, not getting friendly service from health facility, and long waiting times were frequently mentioned factors for not continuing PMTCT service utilization. This is in line with findings of other study done in Ethiopia revealed that health workers are not happy to handle deliveries for women who are known HIV positive due to fear of accidental infection [22]. Other studies have also had similar findings [17, 23, 28-31].

In our study, the main reasons for women not taking PMTCT service mentioned by health workers were disclosure of the result, fear of partner, drug side effect, drug supply interruption, trained health care professional not giving attention to the work and work load. Other studies have also had similar findings [18, 19, 28].

The most frequently mentioned promoting factors related to the community was good willingness to disclosure of HIV status. This is in line with other studies [31]. Although stigma affects women to disclose their HIV status, positive disclosure experiences can help women to get support from family members and partners. This further helps women to overcome barriers to PMTCT service utilization.

Frequently mentioned promoting factors related to individual and family were partner support, disclosure of HIV status, good support from family and peers, previous history of getting HIV free child, healthy feeling, and good awareness on HIV transmission. Continuous Partner support to the women for those who had disclosed their HIV status was reported to the most frequently mentioned promoting factors for PMTCT service utilization. This support can be like remind to take their drugs and counseling. The findings described in this study were similar to those reported in other studies in Zimbabwe [21] partner, community and health worker support were facilitators to PMTCT service utilization. Other study also had similar findings [31]. This emphasizes the great influence that partner have made on the decisions of pregnant women to use PMTCT services.

The findings of this study revealed that mother support groups are facilitator of PMTCT service utilization. The findings also similar with other studies in Malawi reported that promotion of peer counseling schemes for PMTCT [30]. This indicated that mother support group served as peer educators in health facilities to provide education and psychosocial support to HIV positive pregnant and lactating women promotes PMTCT service utilization.

The limitations associated with this study includes: As this was a qualitative study, the results are not generalized to the population at large. Social desirability bias cannot be avoided as the study topic dealt with sensitive issues. This could potentially alter participants’ responses, causing them to provide answers that conform to socially accepted norms. To reduce this bias, highly trained interviewers familiar with qualitative interview techniques were used. Generally, we think that these limitations do not significantly compromise the validity of our study.

## Conclusion

The study has identified numerous facilitators and barriers to PMTCT service utilization which can help in the rollout of PMTCT care among HIV positive women. The major barriers were related community factor, individual and family related factors and health institution factors. Stigma, discrimination, limited knowledge on HIV, lack of partner support, fear of divorce, lack of family support not getting friendly service from health facility, and taking time to get service were major barriers to PMTCT service utilization.

Continuous partner support, previous history of getting HIV free child, good health worker support, good awareness on PMTCT, availability of drugs, and discussion during coffee ceremony with mother support group were the major facilitators for PMTCT service utilization. Strengthening community awareness on HIV, engaging male involvement in PMTCT care and getting friendly service were the key determinants for the better PMTCT service utilization. Future research should be done on PMTCT service provision and utilization problems.

## List of abbreviation

ART: Antiretroviral Therapy
FGD: Focus group discussion
HIV: Human Immunodeficiency Virus
IDI: In depth interview
MTCT: Mother-to-Child Transmission
PMTCT: Prevention of Mother-to-Child Transmission.

## Competing interest

The authors report no conflicts of interest in this work

## Funding

Not applicable

## Authors’ contribution

Z.Z., Y.M., K.Y. and G.A. designed the research study. Z.Z., Y.M., K.Y. and G.A. performed the research. Z.Z. analyzed the data. Z.Z. wrote the draft paper. Z.Z., Y.M., K.Y. and G.A. critically review the paper. All authors approved the final manuscript and agree to be accountable for all aspects of the work.

## Acknowledgments

The authors would like to thank Mr. Awraris Hailu, for his critical review of draft versions of this manuscript and assisting in using ATLAs software. We would like to thank Dr. Mekuriaw Alemayehu for assistance in using ATLAs software. We would also like to acknowledge facilitators and study participants.

